# The African wolf is a missing link in the wolf-like canid phylogeny

**DOI:** 10.1101/017996

**Authors:** Eli K. Rueness, Pål Trosvik, Anagaw Atickem, Claudio Sillero-Zubiri, Emiliano Trucchi

**Affiliations:** Centre for Ecological and Evolutionary Synthesis (CEES), Dept. of Biosciences, University of Oslo, P.O. Box 1066 Blindern, N-0316 Oslo, Norway; Wildlife Conservation Research Unit, The Recanati-Kaplan Centre, Tubney House, Zoology, University of Oxford, Tubney OX13 5QL, UK; Department of Botany and Biodiversity Research, University of Vienna, Rennweg 14, 1030, Vienna, Austria

**Keywords:** Wolves, genome-wide phylogeny, canid evolution

## Abstract

Here we present the first genomic data for the African wolf (*Canis aureus lupaster*) and conclusively demonstrate that it is a unique taxon and not a hybrid between other canids. These animals are commonly misclassified as golden jackals (*Canis aureus*) and have never been included in any large-scale studies of canid diversity and biogeography, or in investigations of the early stages of dog domestication. Applying massive Restriction Site Associated DNA (RAD) sequencing, 110481 polymorphic sites across the genome of 7 individuals of African wolf were aligned and compared with other wolf-like canids (golden jackal, Holarctic grey wolf, Ethiopian wolf, side-striped jackal and domestic dog). Analyses of this extensive sequence dataset (ca. 8.5Mb) show conclusively that the African wolves represent a distinct taxon more closely related to the Holarctic grey wolf than to the golden jackal. Our results strongly indicate that the distribution of the golden jackal needs to be re-evaluated and point towards alternative hypotheses for the evolution of the rare and endemic Ethiopian wolf (*Canis simensis*). Furthermore, the extension of the grey wolf phylogeny and distribution opens new possible scenarios for the timing and location of dog domestication.

## 1. Introduction

Wolf-like canids (genus *Canis*) form a widely distributed group of mammalian carnivores encompassing animals popularly termed wolves, jackals and coyotes. Their common ancestor has been suggested to have originated in Africa about 4 mill years ago [1]. Insights into the biogeography and phylogeny of several of the members of the group have, however, been hampered by difficulties with taxonomic classification and the existence of cryptic species.

Within the larger group of wolf-like canids grey wolves form a species-complex which, in addition to the widely distributed Holarctic grey wolf (HGW, *Canis lupus*), includes the domestic dog (*Canis lupus familiaris*) and three more ancient mitochondrial DNA (mtDNA) lineages that have recently been discovered in Asia [2] and in Africa [3].

Notions about the presence of grey wolves on the African continent, in particular in Egypt, go all the way back to the ancient Greek historian Herodotus in the 5^th^ century BC and can also be found in the writings of famous naturalists of the classical antiquity such as Aristotle and Pliny the Elder [4]. In the mid-19^th^ century the resemblance of North African jackals to grey wolves inspired the systematic name *Canis lupaster* to emphasize their status as a wolf-like species different from the more widely distributed golden jackals (*C. aureus*) of Africa and Eurasia [5]. The taxonomic status of *C. a. lupaster* has since been vividly debated [6]. Based on morphological traits (size and craniological measures) the classification has alternated between *i)* unique species, *ii)* subspecies of grey wolf and *iii)* subspecies of golden jackal. Presently *C. a. lupaster* (Hemprich and Ehrenberg 1833) is classified as golden jackal and thus categorized as a species of least concern by the IUCN [8]. The first genetic evidence connecting *C*. *a. lupaster* to the grey-wolf species complex was published in 2011 [3], and its distribution range was soon confirmed to be much larger than anticipated [9]. All mtDNA sequences published for African canids with a golden jackal phenotype have had a distinctive wolf-like haplotype [3,9,10]. These results open up the possibility that animals previously thought to be golden jackals are in fact African wolves. However, since all of these studies were based on maternally inherited mtDNA, and hybridization is common among wolf-like canids (e.g [11,12]), the possibility of a hybrid origin of the African wolf could not be excluded, emphasizing the need for thorough genomic studies.

In spite of numerous studies the timing and location of canid domestication remains arduously disputed. Recently published whole genome sequence data indicate that HGWs and domestic dogs are monophyletic sister clades [10]. Thus, inclusion of the full taxonomic diversity within the grey wolf species-complex may shed new light on arguably one of the most ancient and important symbiotic relationships between humans and domestic mammals.

Here, we provide the first genome-wide sequence data for African wolves and address questions pertaining to its origin and phylogenetic relationship to HGWs, the golden jackal and other wolf-like canids, including the rarest canid in the world the Ethiopian wolf (*Canis simensis*) [13], and the domestic dog. Since all our African wolf individuals were sampled in Ethiopia we will hence refer to the endemic Ethiopian wolf as *C. simensis,* in order to avoid confusion.

Particularly we aim to address the following questions:

1. Is the African wolf a hybrid between other wolf-like canids?
2. Is the African wolf a species separate from golden jackals?
3. What is the relationship between the African wolf and *C. simensis*?
4. Could the African wolf have been a key to dog domestication?

## Results

### RAD sequence data

Reduced-complexity methodologies in genome typing, such as Restriction site Associated DNA sequencing (RADseq: [14]), facilitate screening of hundreds of thousands of loci across multiple individuals in non-model species (e.g. [15]). We used the Illumina HiSeq platform to RAD-sequence 16 canid tissue samples: 7 African wolves (Ethiopia), 2 HGWs (Norway), 2 *C. simensis*, 2 golden jackals (Serbia) and 3 side-striped jackals (*Canis adustus*, Ethiopia), see Table S1 for details. The resulting sequence data were processed and aligned using the Stacks pipeline [16], resulting in 90211 marker loci containing a total of 110481 polymorphic sites (i.e. aligned positions with at least two base variants). The loci were subsequently aligned to the domestic dog reference genome and 79752 corresponding sequences were retained. The mean number of polymorphic sites in our individuals was 28342 (± 16777) and the number of polymorphic sites in each individual is presented in Figure S1. The large variation in numbers of polymorphic sites is due to variability in Illumina read numbers among individuals.

### The African wolf is not a hybrid species

Subsets of the loci from the unreferenced and referenced datasets (the referenced data had been aligned to the dog genome prior to variant calling) were selected by two different criteria (see Methods below) and used for Bayesian cluster analysis in order to investigate the genetic structuring among the five species. The most likely number of clusters in the data (*K*) was shown to be five (values of *K* between 2 and 8 were tested; Figure S2). When allowing five clusters all of the individuals were assigned to their putative species with a probability of > 0.99. These results strongly suggest that the African wolf should be considered a unique species that has not resulted from a historical hybridization event. Interspecific hybridization is known to occur among wolf-like canids in the wild [12,17] and we ran the admixture model five generations back allowing the individuals to have partial ancestry in each cluster. None of the individuals had a higher probability than 0.01 of having recent hybrid ancestors.

### The African wolf is distinct from the golden jackal

The calibrated mtDNA phylogeny of the canid species represented in this study (Figure 1A) shows that the African wolf falls within the grey wolf species-complex, and the presence of canids with the African wolf haplotype has been confirmed in several countries in both East –and West Africa. (Figure 1B). We produced a neighbor-joining network of the17 canid individuals (Figure 2) based on all of the sites that were variable in at least one individual from the full RADseq dataset (including loci with missing data across samples). Side-striped jackals, *C. simensis* and golden jackals all form clearly diverged clusters. The genetic diversity among the seven individuals of African wolf is high, but they form a group along with HGWs and the domestic dog, which is clearly distinguished from the golden jackals.

**Figure 1:**
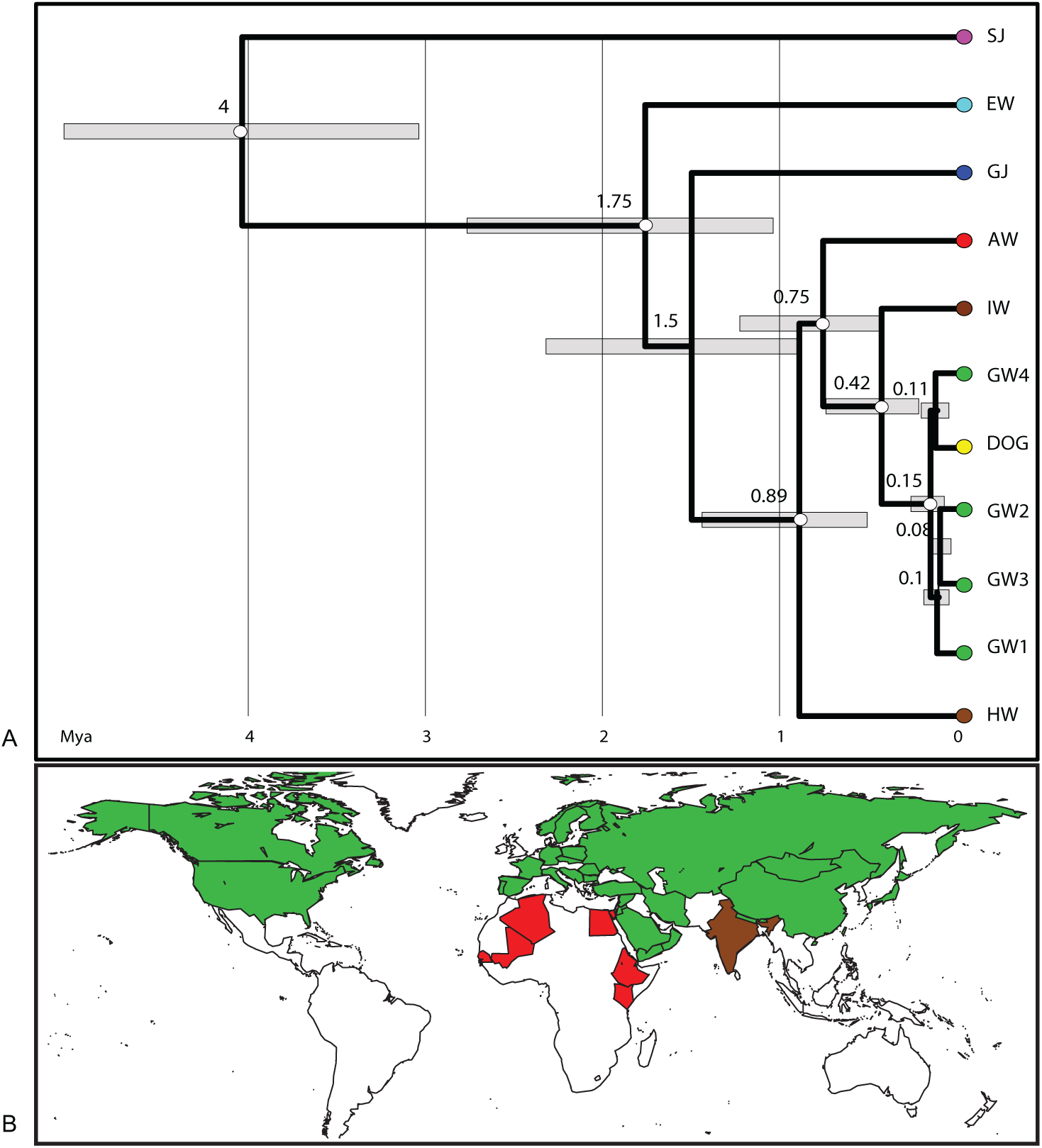
mtDNA phylogeny and geographic distribution of the grey wolf species-complex. A) The calibrated phylogeny is based on 1119 bp of the Cyt*b* gene and 316 bp of the control region. The grey wolf species-complex includes the Holarctic grey wolf (HGW; from Canada (GW2), China (GW3) Norway (GW1) and Saudi-Arabia (GW4)), the Indian wolf (IW), the Himalayan wolf (HW) and the African wolf (AW). In addition the Ethiopian wolf (*C. simensis*; EW), the golden jackal (GJ) and the side-striped jackal (SJ) are included. Bayesian posterior probabilities above 99% are shown at the branching points as white circles. The average estimated age (in million years) of each node is shown with the 95% HPD as grey bars. B) Global distribution (by countries) of the grey wolf species-complex indicated by color shading: the HGW = green, the Indian and Himalayan wolves = brown, and the African wolf = red. Only countries where the presence of the African wolf has been confirmed by genetic analysis (mtDNA) are shaded.

**Figure 2:**
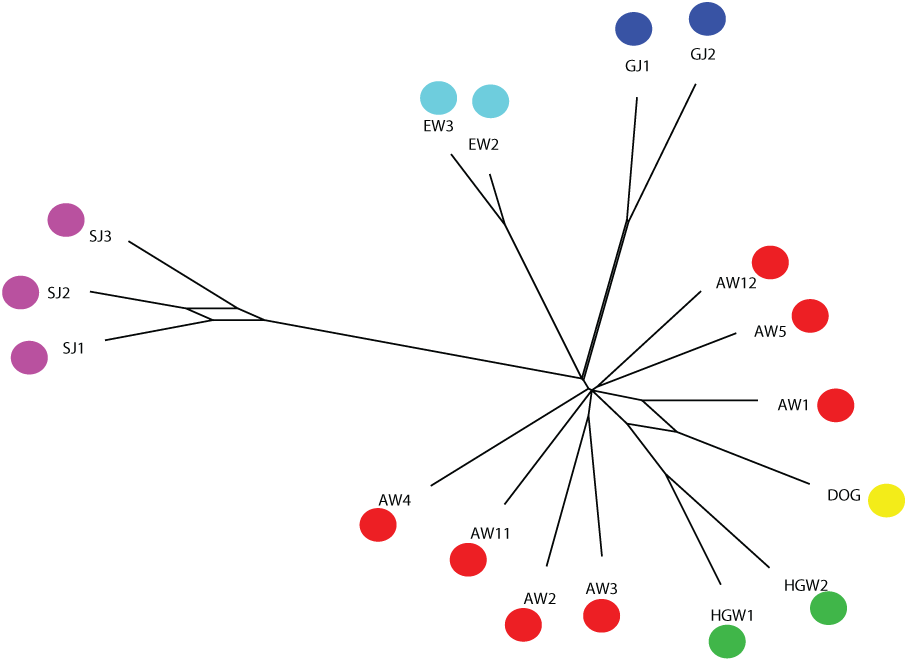
Distance network based on the full set of SNPs. Consensus network based on neighbor-net analysis of 11,0481 polymorphic sites, displaying the phylogenetic relationships among all the studied individuals. All of the major branches had 100% bootstrap support, while the splits between some of the African wolves as well as between the two individuals of *C. simensis* were less confident. The following abbreviations and color coding were used: AW (red) = African wolf, HGW (green) = Holarctic grey wolf, GJ (dark blue) = golden jackal, EW (light blue) = Ethiopian wolf (*C. simensis*), SJ (purple) = side-striped jackal, DOG (yellow) = domestic dog.

As the amount of missing data was large for many of the sequenced samples we merged polymorphic sites within each species and then removed all sites with missing data for one or more species (see Methods below). This produced an alignment of 8001 polymorphic sites with each site represented in at least one of the individuals within each of the species. Figure S3 shows the number of private SNPs observed per species (mean = 957) with the side-striped jackal clearly being the most divergent (2821 private SNPs). Relative distances among species (Figure S4) followed the same pattern as the network that included loci with missing data (Figure 2).

Phylogenetic relationships among the species reconstructed as a ML tree (Figure 3A) based on 768 concatenated loci (72960 bp with 1388 polymorphic sites) showed essentially the same topology as the mtDNA tree (Figure 1A) and the networks based on the full or merged data sets (Figure 2, Figure S4). The node connecting the African wolf to the HGW and dog was strongly supported with a bootstrap value of 100%. The side-striped jackal was the most basal species, consistent with earlier findings [1]. A calibrated species tree was reconstructed through a multi-locus Bayesian approach using random selections of 30 RAD loci with 4 to 6 SNPs and also including a partial sequence of the Cytochrome *b* (Cyt*b*) gene. In this analysis, the timing of the split between the golden jackal and the wolf ancestor was 1.35 mya (95% HPD: 0.95-1.94 mya; Figure 3), which is consistent with previous estimates based on both fossil and genomic data [1,18]. When using only mtDNA for phylogenetic calibration the split was estimated at 1.5 Mya, (95% HPD: 0.88-2.32 Mya; Figure 1A). The main difference between the phylogenies based on genomic and mtDNA data is the position of *C. simensis*. However, this node was not highly supported in any of the analyses.

**Figure 3:**
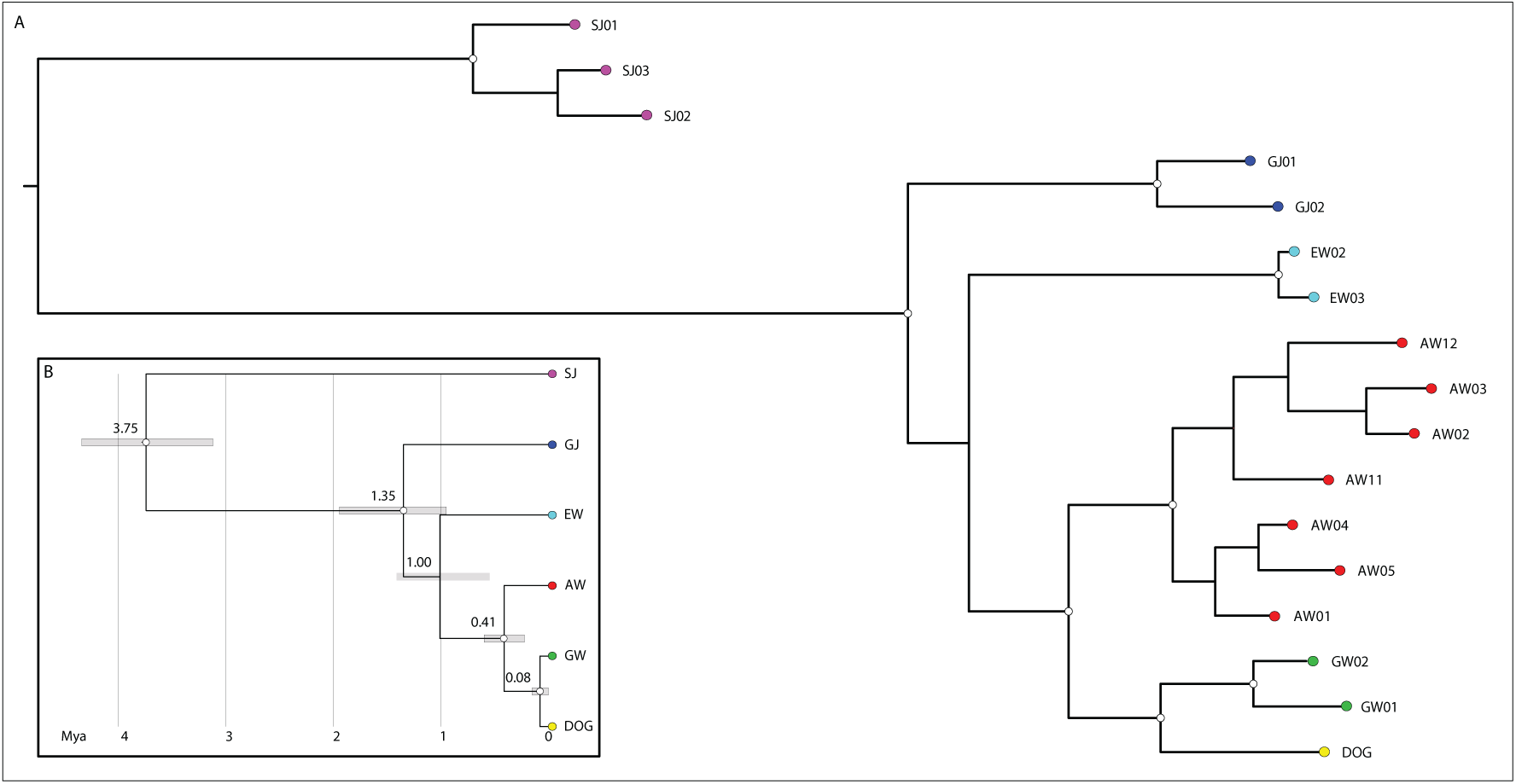
ML tree and divergence time estimates based on Bayesian species tree analysis. In the main figure (A) a ML tree was based on 72,960 bp with 1,388 polymorphic sites from 768 concatenated loci. The inset (B) shows the species tree inferred by the *beast algorithm based on a random set of 30 RAD loci and mtDNA Cyt*b* partial sequence. The average estimated age (in million years) of each node is shown with the 95% HPD as grey bars. Bootstrap support of 100% (A) and bayesian posterior probabilities above 99% (B) are indicated by white circles on the nodes. Tip labelling and color coding of species are the same as in Figure 2.

The phylogenetic analyses presented here clearly demonstrate that African canids currently classified as golden jackals based on phenotypic criteria are genetically distinct from golden jackals from Eurasia.

### An African origin of the Ethiopian wolf (Canis simensis)?

The evolutionary history of Africa’s most endangered carnivore, *C. simensis*, is enigmatic, but it has been hypothesized that a recent ancestor migrated from Eurasia into the Horn of Africa [19]. Based on low diversity in the mtDNA control region (CR) of *C. simensis* their divergence from a HGW/coyote (*Canis latrans*) cluster was estimated at around 100,000 years ago [20]. However, due to extremely low population sizes associated with the patchy availability of Afroalpine habitats (only some 500 adult individuals currently remain in six populations [21]) this estimate should be treated with some caution.

Our analyses based on nuclear and mitochondrial data indicate that *C. simensis* form a sister clade to African wolves, grey wolves and domestic dogs. Thus, rather than postulating a Eurasian emigration event a more parsimonious hypothesis would be that *C. simensis* and African wolves shared a common ancestor until approximately 1 mya ago (95% HPD: 0.55-1.41 mya), when *C. simensis* diverged and evolved into specialist Afroalpine rodent hunters [22] without having left Africa.

### Are-evaluation of the location of dog domestication

Our phylogenetic analysis (Figure 3) suggests that the AW shares a common ancestor with the HGW and the dog about 0.41 mya (95% HDP: 0.22-0.60 mya; 0.75 Myr according to mtDNA data only). We estimated the timing of the split between HGWs and domestic dogs at 80,000 years ago (95% HPD: 0.0002-0.151 mya), while earlier estimates based on genetic data ranges between 11,000 to 135,000 years ago [10,23,24].

In the network, the node connecting the domestic dog, the HGWs and one of the African wolves there is a reticulum (Figure 2), indicating that this particular connection is poorly resolved. This could be explained by data deficiency, as the African wolf individual in question (AW1) has a relatively low number of sampled polymorphic sites (Figure S1). Alternatively, it could indicate introgressed dog genes in this particular individual, or that African wolves have contributed to the dog’s genomic history.

## Discussion

RADseq is a convenient way of sampling large numbers of loci from multiple individuals. One potential caveat of this technique is uneven sampling between individuals, both in terms of quantity and distribution [25]. In our data differences are considerable and three African wolves, as well as *one C. simensis* and one side-striped jackal, are somewhat under-sampled (Figure S1). However, the results presented here are still based on large amounts of sequence data and robust analytical approaches.

In hybrid species one would expect discordance between mitochondrial and nuclear lines due to introgression of nuclear genetic material. The topologies of the phylogenetic trees presented here were consistent with respect to the position of the African wolf regardless of whether they were based on mtDNA (Figure 1A) or genome-wide data (Figure 3), and Bayesian admixture analysis did not detect any signal of a hybrid origin in our seven African wolf individuals. This does not exclude the possibility that African wolves are involved in hybridization events with closely related sympatric species, as inter-specific mating is not uncommon in wolf-like canids [26], including *C. simensis* [20]. We are currently extending the analysis to African wolves from other parts of the continent to explore the possibility of hybridization further and to study the patterns of intra-specific variability.

According to current perception animals known as golden jackals form a single species and are distributed throughout Northern and Central Africa and in Southern Eurasia from the Balkans to South China [8]. Recent studies [3,9] have indicated that this wide distribution range may be grossly exaggerated due to misclassification of African wolves. Out of 24 published partial mtDNA sequences from six countries spanning the presumed African distribution range of this putative species, not a single one is similar to the golden jackal haplotype from Eurasia. Thus, the presence of golden jackals in Africa is questionable, or at the very least their distribution range has been severely overestimated.

*C. simensis* is sympatric with African wolves, normally referred to as ‘common jackals’ in the Ethiopian highlands. The reigning hypothesis regarding the origin of Ethiopian wolves is strictly conceptual, as no fossilized animals resembling *C. simensis* have been found to date. Our data open up the possibility for an African origin for the species, from a common ancestor shared with African wolves. Full genome sequences of both species should be compared in order to gain further insight into this question.

Similarly, the presence of ‘grey wolves’ on the African continent has implications for questions pertaining to dog domestication. In spite of disagreement between researchers as to the timing, location and number of events, it is generally thought that the HGW domestication took place somewhere in Eurasia. A recent genomic study by Freedman *et al.* [10] suggested that modern day dog lineages contain genetic variation not found in HGW from any of the suggested source populations and concludes that wolves and dogs are sister taxa sharing a common ancestor. No previous genomic study addressing domestication has included Indian, Himalayan or African wolves, although the high mtDNA diversity detected in African and Asian village dogs [27] suggests that dogs have a long evolutionary history in these areas. We therefore suggest that the full phylogenomic diversity and geographical range of wolves and dogs may provide valuable clues to the history of man’s best friend.

## Methods

### Sampling DNA extraction and mtDNA sequencing

Tissue samples from seven African wolves, and three side-striped jackals were collected from carcasses in the field. The work was permitted by the Ethiopian wildlife conservation authority (EWCA), and since no live animals were involved no ethical approval was required. The two samples of *C. simensis* were collected from live animals trapped as part a bilateral agreement between the University of Oxford and the EWCA. This research project follows the guidelines of, and was approved by, the Zoology Ethical Review Committee (University of Oxford), Ref ZERC040905. We follow the ASM standard [28]. Two tissue samples from legally hunted Holarctic gray wolves were collected by the Norwegian Institute for Nature Research (NINA) with permission from the State Nature Inspectorate. Samples from two golden jackals were collected for a previous study by Zachos *et al.* [29]. See Table S1 for details. DNA was extracted using the DNeasy Blood & Tissue kit (Qiagen) according to the manufacturer’s instructions. DNA quantity and quality were ascertained by fluorometric-based measurement on a Qubit® 2.0 Fluorometer (Life Technologies) and by gel electrophoresis.

The Cyt*b* (1119 bp) gene was amplified in two fragments by the primer-pairs cytb1 – cytb2 [30] and L15162 – H1595 [31]. A 316 fragment of the CR was amplified by the primers Thr-L 15926 and DL-H 16340 [32]. Sequencing and electrophoresis were performed on an ABI 3730 PRISM machine (Applied Biosystems). The mtDNA haplotype sequences that are not previously published have been uploaded to GenBank and assigned the following accession numbers: *C. simensis* (Cyt*b*: KP874954), Side-striped jackal (Cyt*b*: KP874955, CR: KP874956).

### mtDNA phylogeny

*MEGA* version 5.2.2 [33] was used to manually align our sequences to the corresponding sequence fragments for the domestic dog (NC_002008), African wolf (Cyt*b*: HQ845258, CR: HQ845259), Himalayan wolf (Cyt*b*: AY291431, CR: AY333740), Indian wolf (Cyt*b*: AY291432, CR: AY289984), *C. simensis* (CR: HQ845261) and HGWs from Saudi Arabia (DQ480506), Canada (DQ480508), China, (KC461238) and Sweden/Finland (Cyt*b*: KF661052, CR: KF723521). A calibrated phylogeny was estimated with the BEAST package version 1.8 [35,36] using the following settings: the two partitions, Cyt*b* and CR were unlinked in respect to substitution model and clock-rate model but linked in respect to tree model as they are part of the same chromosome. MrModeltest 2.3 [37] was used to find the optimal substitution models as defined by the Akaike Information Criterion (AIC). The selected substitution models were HKY+I+G for CR and HKY for Cyt*b*. In the latter, a 3-partitions substitution model was also used to accommodate a different rate for each codon position. A log-normal relaxed clock model was chosen for each partition with the mean value estimated from the data. As we only included samples from different species/populations, a Birth-Death model was set as tree model. We then constrained the root of our tree (split between *Canis adustus* and *Canis lupus*) on the basis of the estimate of 4.3 Mya (credibility region between 3.4 and 5.5 Mya) provided by Perini *et al.* [1]. A normal distribution with mean 4.3 Myr and stdev 0.5 Myr was set for the time since most recent common ancestor (tmrca) of all taxa included in our tree. Three replicates were run for 10,000,000 generations and convergence of parameters was checked on Tracer 1.5 (http://tree.bio.ed.ac.uk/software/tracer/) by examination of effective sample size (ESS) values. Final summary trees were produced in TreeAnnotator ver 1.7. and viewed in FigTree 1.4 (http://tree.bio.ed.ac.uk/software/figtree/).

### RAD library preparation and sequencing

The following RADseq protocol was adopted from that outlined in [14]: *i*) approximately 50ng of genomic DNA per sample were digested with the restriction enzyme *Sbf*I (NEB); *ii*) each sample was then ligated to a unique barcoded P1 adapter (see Table S1) prior to pooling in a single library. The library was then sheared by sonication, and gel electrophoresis of small library aliquots were run after the first 5 cycles (30″ ON – 30″ OFF) and then every 1-2 cycles of sonication; *iii*) the target size range fraction (300-500 bp) was achieved after 8 cycles of sonication and was then selected by gel electrophoresis and manual excision; *iv*) before size selection on the gel, sonicated libraries were concentrated to 25 μl by DNA capture on magnetic beads (beads solution:DNA = 0.8:1), thus further reducing the carry-over of non-ligated P1 adapters; *v*) capture on magnetic beads using the same beads:DNA ratio (0.8:1) was then employed in all following purification steps (after blunt-end repairing, poly-A tailing, P2 adapter ligation and library enrichment by PCR); *vi*) PCR amplification (23 cycles) was performed in 8 × 12.5 μl aliquots pooled after the amplification in order to reduce amplification bias on few loci due to random drift; *vii*) the library was then quantified by a fluorometric-based method (Qubit® 2.0, Life Technologies) and molarity was checked on an Agilent Bioanalyzer chip (Invitrogen). A final volume of 20 μl with a DNA concentration of 45 ng/μl was sequenced using the ILLUMINA HiSeq2000 platform at the Norwegian Sequencing Centre, University of Oslo.

### Quality filtering and SNP calling

The sequencing resulted in 179,146,995 reads of 100bp. Raw reads were processed on the ABEL computing cluster (University of Oslo) using the scripts included in the Stacks package, a software pipeline for building loci from short-read sequences [16]. The sequencing resulted in 179,146,995 reads of 100bp. Raw reads were quality filtered and de-multiplexed using “process_radtags.pl” with default parameter settings. Catalogue building using “denovo_map.pl” was carried out with the following settings: m (minimum number of identical, raw reads (i.e. coverage) required to create a stack/locus) = 3, n (the number of mismatches allowed between loci when building the catalogue) = 7, M (the number of mismatches allowed between loci when processing a single individual) = 5. A table including all loci matching the 16 sequenced individuals was built using “export_sql.pl”. We also aligned the filtered and de-multiplexed sequences to the dog reference genome (see below) using Bowtie2 [38] and built an alternative referenced loci catalogue from the resulting SAM files using “ref_map.pl”.

Several different settings of read trimming parameters, quality thresholds and mismatches were tested when building the individual and the population catalogues to check for consistency of the results.

Loci with more than two variants in two individual were discarded. For loci that were heterozygous in an individual one variant was randomly selected. These processing steps were carried out using custom R-scripts [38].

### Cluster analysis

STRUCTURE version 2.3.4 [39] was used to explore whether all individuals were assigned to their putative species. We analysed two set of loci selected by different criteria. *i*) 1051 polymorphic loci matching more than 10 samples and with a maximum of 6 SNPs were extracted from the catalogue built in Stacks using the denovo_map.pl command. Using custom python scripts, these loci were further filtered in order to exclude loci with more than 2 alleles occurring in one or more samples and loci with at least one SNP in the last 10 base pairs. We decided to apply this strict filtering criterion after checking the SNP distribution along the loci length (95 base pairs) and verifying a higher than average occurrence of SNPs in the last 10 base pairs. Our dataset was then reduced to 768 loci with 1 to 6 SNPs. *ii*) The “populations” program in Stacks were used to select 2849 referenced loci with the following settings: r (minimum percentage of individuals in a population required to process a locus for that population) = 0.25, p (minimum number of populations a locus must be present in to process a locus) = 2, and m (minimum stack depth required for individuals at a locus) = 6. To determine the most likely number of clusters in the data (*K*) the length of the burn-in was we 10,000, and we ran 100,000 MCMC iterations for numbers of *K* from 2 to 8. We ran the program 3 times per value of *K* and then calculated the differences in likelihood, delta *K* [40] applying the Structure Harvester [41] to determine the optimal number of clusters. To test for recent events of migration we ran the ancestry model using the information of population origin for each individual as priors. The migration prior (rate of migration allowed per generation) was set to 0.05 (values in the range of 0.001—0.1 are suggested by Pritchard *et al.* [39] and gensback (number of generations back in time where migration is assumed) was set to 5.

### Alignment to the domestic dog reference genome

The domestic dog genome assembly CanFam3.1 was downloaded from the Illumina iGenomes collection (http://support.illumina.com/sequencing/sequencing_software/igenome.ilmn). RADSeq markers were aligned to the full reference genome using BLAST+ [42] version 2.2.26 for retrieval of coordinates of corresponding sequences in the domestic dog genome. For each marker the coordinates were taken from the top scoring BLAST hit with at least 85% sequence identity. 79752 full length marker sequences were found in the dog reference genome (from a total of 110481, >80%), and these were extracted using genomic coordinates and a custom R-script [38]. Domestic dog marker sequences were inserted into the main alignment and SNPs were extracted and concatenated using a custom R-script [38].

### Neighbor Joining Network analysis

Phylogenetic networks were made with Splitstree 4.13 [43] using the neighbor-net algorithm for agglomerative clustering based on a matrix of genetic distances. The confidence of the branches was tested through bootstrapping (1000 replicates) and a confidence network, displaying the splits that appeared in at least 95% of the input trees, were constructed.

SNPs were merged within each species in the following way: if the SNP was found in one individual within a species that nucleotide was taken to represent the species in that site. If the SNP was found in more than one individual a nucleotide was sampled randomly from those individuals. Finally, SNPs that were not represented in all 6 species were removed from the alignment, resulting in a total of 8001 polymorphic sites.

### Phylogenetic analysis using the genomic dataset

768 RAD loci were selected as described above (cluster analysis) and concatenated to a single sequence per sample encoding the heterozygosities as ambiguities (IUPAC code) if two alleles were present [44]. The whole sequence of each locus was included in order to get empirical estimates of the base composition and invariant sites. This dataset was then employed in a Maximum-Likelihood tree reconstruction using a GTR+Gamma substitution model and 100 rapid bootstraps in RAXML v.7.2.8 [45]. A subset of 30 RAD loci chosen at random among those with the highest number of SNPs (4 to 6 SNPs) were employed in a multi-species coalescent analysis using the *beast algorithm [46] implemented in BEAST 1.8. Each individual was assigned to its putative species. Cyt*b* sequence data were also included in this analyses. Three runs of 50 million iterations using different random selections of RAD loci were compared. For the genomic data loci were unlinked for site substitution model, clock model and tree prior model; site substitution models were set as a HKY with empirical base frequency; a strict molecular clock was estimated for each marker with a non-informative uniform prior distribution bounded within 0.05 and 0.0005 sub/s/Myr. For the Cyt*b* data the same settings employed in the mtDNA analysis (described above) were applied: a HKY and 3-partitions substitution model and a relaxed clock rate whose ucld.mean was set to 0.025 sub/s/Myr as it was estimated in the previous analysis. As in the mtDNA analysis, we constrained the root of the tree (i.e. the split between *Canis adustus* and *Canis lupus*) using a normal distribution with mean 4.3 Myr and stdev 0.5 Myr (see above for references). The Yule model was selected as tree prior model for the species tree together with a constant population coalescent prior inside each species (Yule - Piecewise constant). Results were visualized and edited in FigTree.

## Acknowledgements

We thank Øystein Flagstad and Frank Zachos for providing samples of Holarctic grey wolves and golden jackals.

---

**Table S1.** Details about the samples analysed.

| # | ID | Locality | Adapter | RAD-Tag |
| --- | --- | --- | --- | --- |
| 1 | AW1 | Ethiopia 1 | P1 | AAAAA |
| 2 | AW2 | Ethiopia 1 | P2 | AACCC |
| 3 | AW3 | Ethiopia 1 | P3 | AAGGG |
| 4 | AW4 | Ethiopia 2 | P40 | GGAAG |
| 5 | AW5 | Ethiopia 3 | P4 | AATTT |
| 6 | AW11 | Ethiopia 4 | P6 | ACCAT |
| 7 | AW12 | Ethiopia 4 | P13 | ATATC |
| 8 | GW1 | Norway | P26 | CGCGC |
| 9 | GW2 | Norway | P27 | CGGCG |
| 10 | GJ1 | Serbia | P19 | CAGTC |
| 11 | GJ2 | Serbia | P25 | CGATA |
| 12 | EW2 | Ethiopia 4 | P16 | ATTAG |
| 13 | EW3 | Ethiopia 4 | P17 | CAACT |
| 14 | SJ1 | Ethiopia 5 | P14 | ATCGA |
| 15 | SJ2 | Ethiopia 6 | P15 | ATGCT |
| 16 | SJ3 | Ethiopia 3 | P5 | ACACG |

Sampling localities in Ethiopia:
1: Arussi, 2: Abyata, 3: Ziway, 4: Bale Mountains
5: Jimma, 6 Addis Ababa

**Figure S1:**
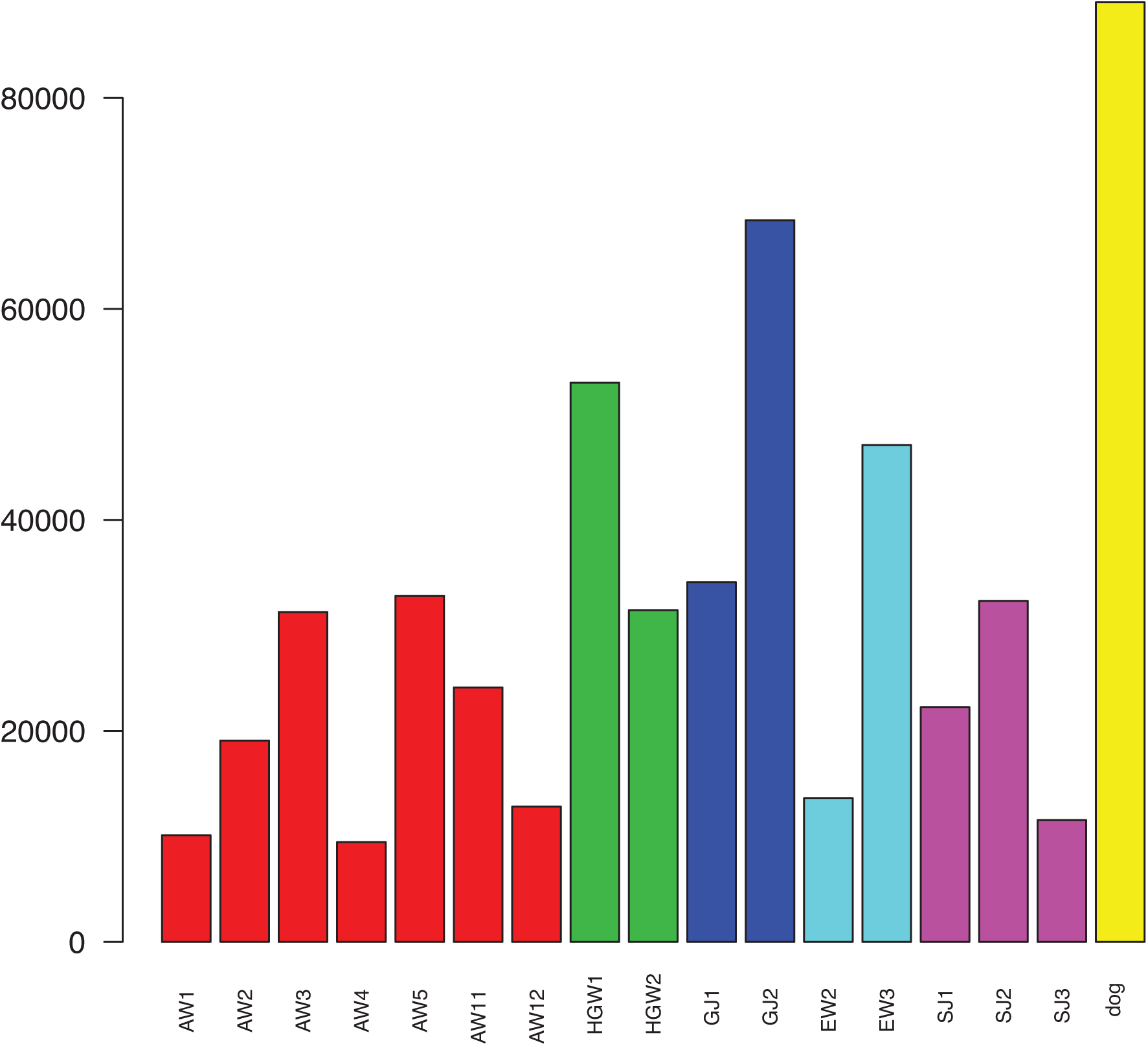
Number of ploymorphic sites per sequenced individual. Individuals are along the x-axis: AW – African wolf (red), HGW – (green), GJ – golden jackal (dark blue), EW – Ethiopian wolf (*C. simensis*; *light* blue), SJ - side-striped jackal (purple). The domestic dog is represented by the yellow bar. The y-axis indicates the number of observed polymorphic sites per individual, and is directly proportional to the amount of sequence data available. The reason for the much higher number of polymorphic sites in the dog is the fact that we had access to the full genome sequence, whereas for all other individuals we only had RAD fragments, resulting in much missing data.

**Figure S2:**
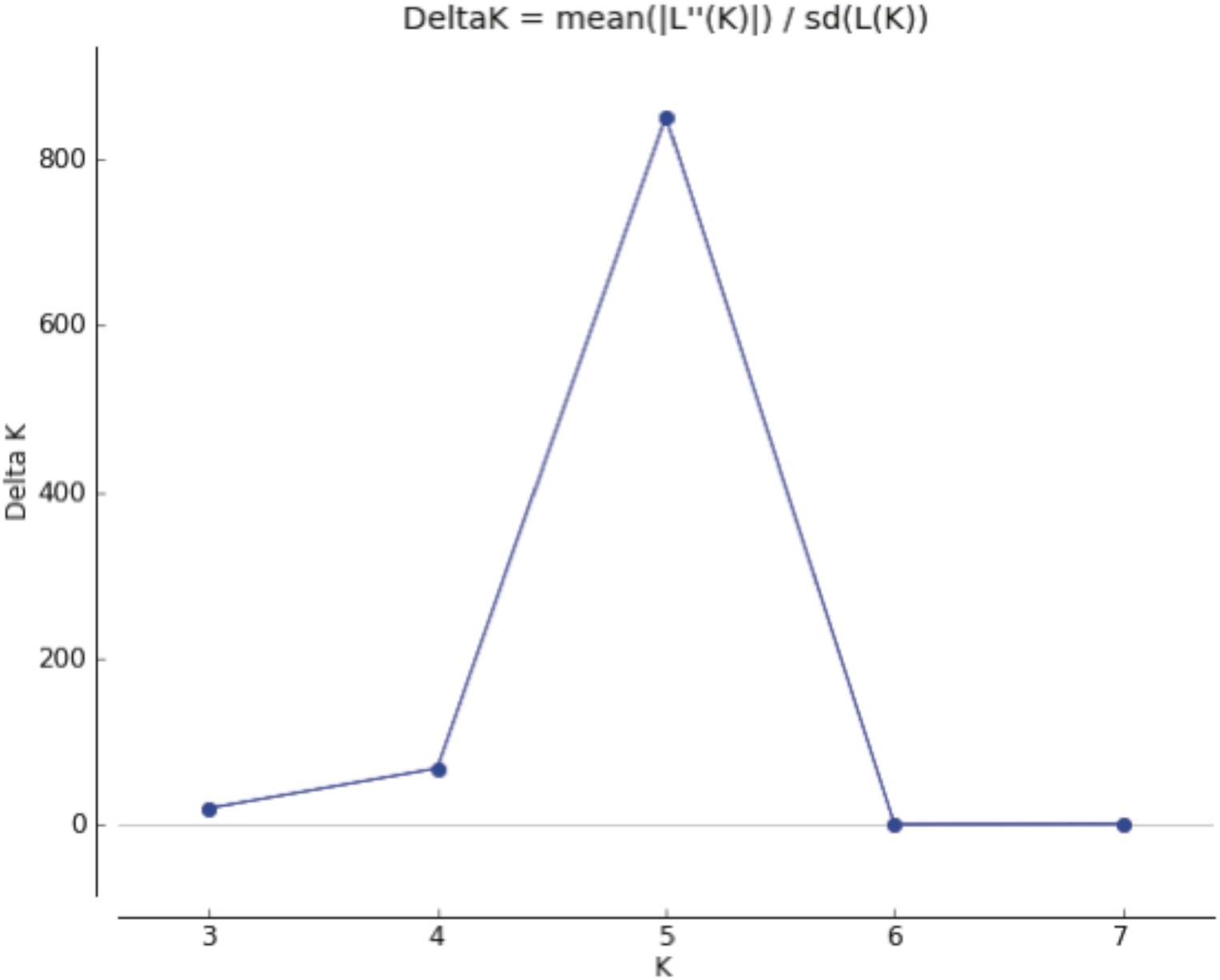
Population assignment analysis. Differences in likelihood for the different numbers of clusters (delta *K*) based on the mean of three runs for each *K.*

**Figure S3:**
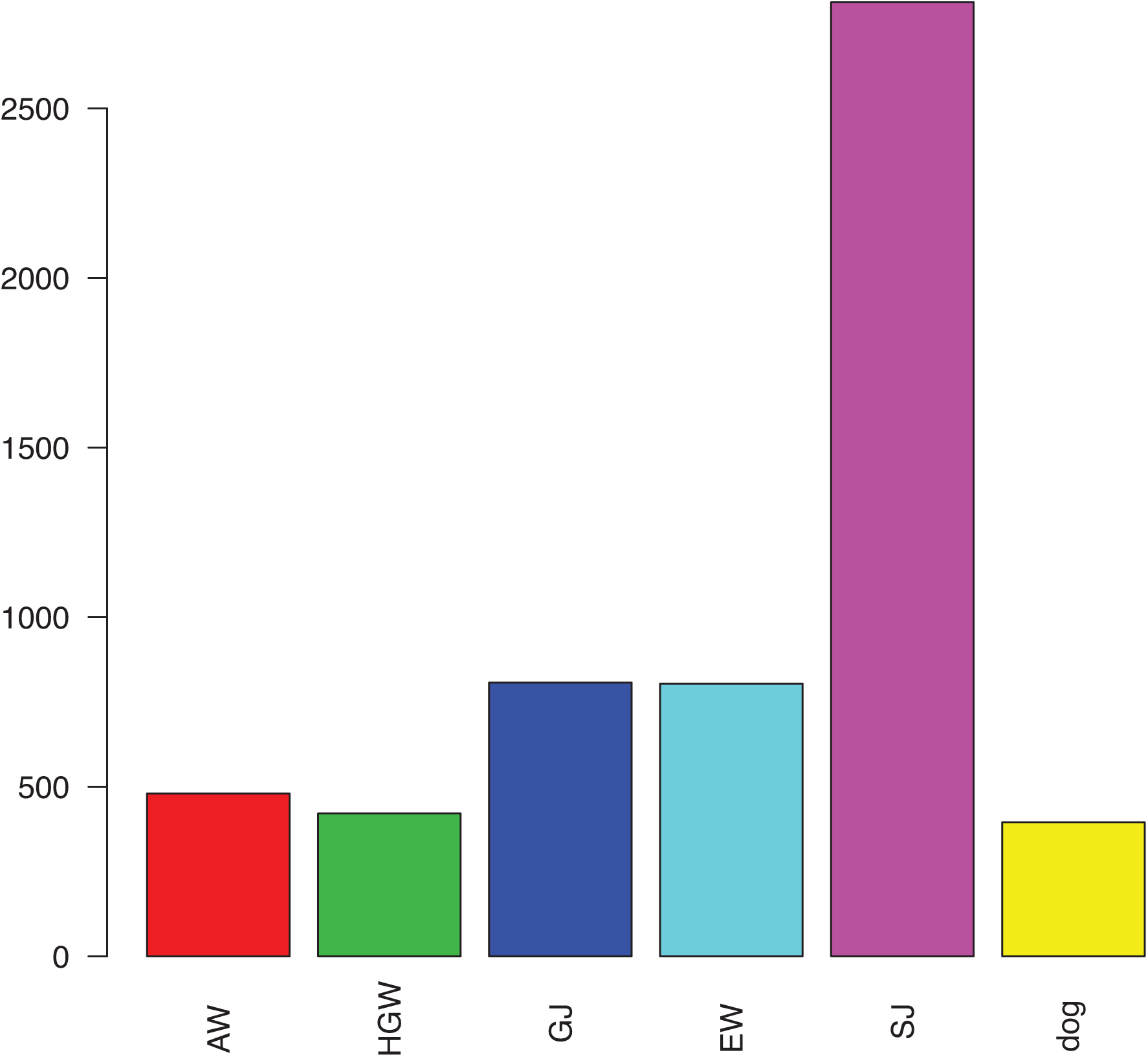
Number of SNPs per sequenced species after data merging. Species are along the x-axis: AW – African wolf (red), GW – HGW (green), GJ – golden jackal (dark blue), EW – Ethiopian wolf (*C. simensis*; light blue), SJ - side-striped jackal (purple). The domestic dog is represented by the yellow bar. The y-axis indicates the number of observed private SNPs.

**Figure S4:**
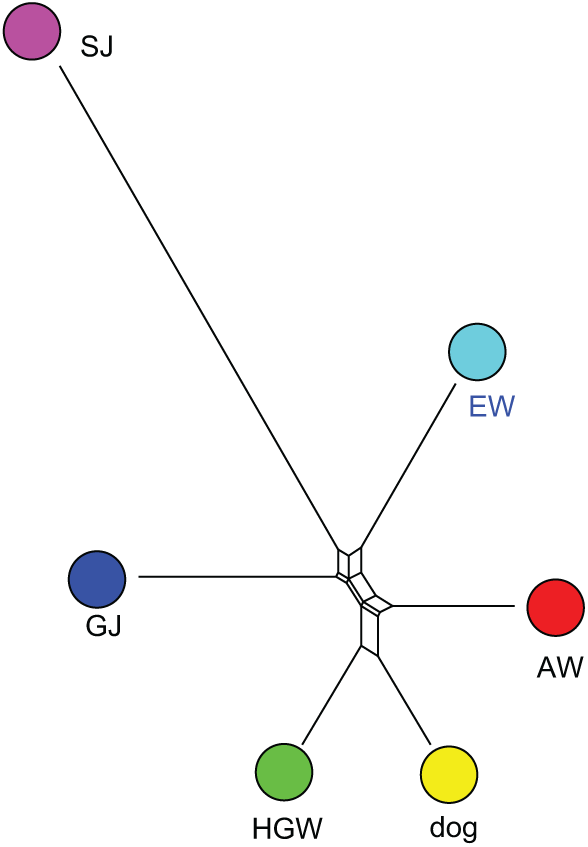
Distance network based on the merged set of SNPs. Consensus network (95% confidence) based on neighbor-net analysis of 8001 SNPs, after data merging, displaying the phylogenetic relationships among the studied species. Labelling and color coding of species as above.

## References

1. Perini FA, Russo CA, Schrago CG (2010) The evolution of South American endemic canids: a history of rapid diversification and morphological parallelism. J Evol Biol 23: 311–322.

2. Sharma DK, Maldonado JE, Jhala YV, Fleischer RC (2004) Ancient wolf lineages in India. Proc Biol Sci 271 Suppl 3: S1–4.

3. Rueness EK, Asmyhr MG, Sillero-Zubiri C, Macdonald DW, Bekele A, et al. (2011) The cryptic African wolf: Canis aureus lupaster is not a golden jackal and is not endemic to Egypt. PLoS One 6: e16385.

4. Larcher, Rennell, Mitford, Schweighæuser (1824) The history of Herodotus. Oxford.

5. Smith CH, Jardine SW (1839) The natural history of dogs: canidae or genus canis of authors; including also the genera hyaena and proteles. Edinburgh: W. H. Lizars.

6. Ferguson WW (1981) The Systematic Position of Canis-Aureus-Lupaster (Carnivora, Canidae) and the Occurrence of Canis-Lupus in North-Africa, Egypt and Sinai. Mammalia 45: 459–465.

7. Wilson DE, Reeder DM (2005) Mammal species of the world : a taxonomic and geographic reference Baltimore, Md: Johns Hopkins University Press.

8. Jhala Y, Moehlman PD (2008) Canis aureus. In: IUCN 2013. IUCN Red List of Threatened Species.

9. Gaubert P, Bloch C, Benyacoub S, Abdelhamid A, Pagani P, et al. (2012) Reviving the African wolf Canis lupus lupaster in North and West Africa: a mitochondrial lineage ranging more than 6,000 km wide. PLoS One 7: e42740.

10. Freedman AH, Gronau I, Schweizer RM, Ortega-Del Vecchyo D, Han E, et al. (2014) Genome sequencing highlights the dynamic early history of dogs. PLoS Genet 10: e1004016.

11. Monzon J, Kays R, Dykhuizen DE (2014) Assessment of coyote-wolf-dog admixture using ancestry-informative diagnostic SNPs. Mol Ecol 23: 182–197.

12. Randi E, Hulva P, Fabbri E, Galaverni M, Galov A, et al. (2014) Multilocus detection of wolf x dog hybridization in italy, and guidelines for marker selection. PLoS One 9: e86409.

13. Marino J (2003) Threatened Ethiopian wolves persist in small isolated Afroalpine enclaves. Oryx 37: 62–71.

14. Baird NA, Etter PD, Atwood TS, Currey MC, Shiver AL, et al. (2008) Rapid SNP discovery and genetic mapping using sequenced RAD markers. PLoS One 3: e3376.

15. Baxter SW, Davey JW, Johnston JS, Shelton AM, Heckel DG, et al. (2011) Linkage mapping and comparative genomics using next-generation RAD sequencing of a non-model organism. PLoS One 6: e19315.

16. Catchen J, Hohenlohe PA, Bassham S, Amores A, Cresko WA (2013) Stacks: an analysis tool set for population genomics. Mol Ecol 22: 3124–3140.

17. Hailer F, Leonard JA (2008) Hybridization among three native North American Canis species in a region of natural sympatry. PLoS One 3: e3333.

18. vonHoldt BM, Pollinger JP, Earl DA, Knowles JC, Boyko AR, et al. (2011) A genome-wide perspective on the evolutionary history of enigmatic wolf-like canids. Genome Res 21: 1294–1305.

19. Gottelli D, Sillerozubiri C, Applebaum GD, Roy MS, Girman DJ, et al. (1994) Molecular genetics of the most endangered Canid - the Ethiopian wolf *Canis simensis*. Molecular Ecolology 3: 301–312.

20. Gottelli D, Marino J, Sillero-Zubiri C, Funk SM (2004) The effect of the last glacial age on speciation and population genetic structure of the endangered Ethiopian wolf (Canis simensis). Mol Ecol 13: 2275–2286.

21. Marino J, Sillero-Zubiri C (2011) Canis simensis. IUCN Red List of Threatened Species. Version 2013.2.

22. Sillero-Zubiri C, Gottelli D (1995) Diet and feeding-behavior of Ethiopian wolves (*Canis simensis*). Journal of Mammalogy 76: 531–541.

23. Savolainen P, Zhang YP, Luo J, Lundeberg J, Leitner T (2002) Genetic evidence for an East Asian origin of domestic dogs. Science 298: 1610–1613.

24. Wayne RK, Geffen E, Girman DJ, Koepfli KP, Lau LM, et al. (1997) Molecular systematics of the Canidae. Systematic Biology 46: 622–653.

25. Davey JW, Cezard T, Fuentes-Utrilla P, Eland C, Gharbi K, et al. (2013) Special features of RAD Sequencing data: implications for genotyping. Mol Ecol 22: 3151–3164.

26. Stronen AV, Tessier N, Jolicoeur H, Paquet PC, Henault M, et al. (2012) Canid hybridization: contemporary evolution in human-modified landscapes. Ecol Evol 2: 2128–2140.

27. Boyko AR, Boyko RH, Boyko CM, Parker HG, Castelhano M, et al. (2009) Complex population structure in African village dogs and its implications for inferring dog domestication history. Proc Natl Acad Sci U S A 106: 13903–13908.

28. Gannon WL, Sikes RS (2007) Guidelines of the American Society of Mammalogists for the use of wild mammals in research. Journal of Mammalogy 88: 809–823.

29. Zachos FE, Cirovic D, Kirschning J, Otto M, Hartl GB, et al. (2009) Genetic variability, differentiation, and founder effect in golden jackals (Canis aureus) from Serbia as revealed by mitochondrial DNA and nuclear microsatellite loci. Biochem Genet 47: 241–250.

30. Janczewski DN, Modi WS, Stephens JC, O’Brien SJ (1995) Molecular evolution of mitochondrial 12S RNA and cytochrome b sequences in the pantherine lineage of Felidae. Mol Biol Evol 12: 690–707.

31. Irwin DM, Kocher TD, Wilson AC (1991) Evolution of the cytochrome b gene of mammals. Journal of Molecular Evolution 32: 128–144.

32. Vila C, Savolainen P, Maldonado JE, Amorim IR, Rice JE, et al. (1997) Multiple and ancient origins of the domestic dog. Science 276: 1687–1689.

33. Tamura K, Peterson D, Peterson N, Stecher G, Nei M, et al. (2011) MEGA5: molecular evolutionary genetics analysis using maximum likelihood, evolutionary distance, and maximum parsimony methods. Mol Biol Evol 28: 2731–2739.

34. Drummond AJ, Rambaut A (2007) BEAST: Bayesian evolutionary analysis by sampling trees. BMC Evolutionary Biology 7: 214.

35. Drummond AJ, Suchard MA, Xie D, Rambaut A (2012) Bayesian phylogenetics with BEAUti and the BEAST 1.7. Mol Biol Evol 29: 1969–1973.

36. Nylander JA, Ronquist F, Huelsenbeck JP, Nieves-Aldrey JL (2004) Bayesian phylogenetic analysis of combined data. Systematic Biology 53: 47–67.

37. Langmead B, Salzberg SL (2012) Fast gapped-read alignment with Bowtie 2. Nature Methods 9: 357–359.

38. R Development Core Team (2014). R: A language and environment for statistical computing. R Foundation for Statistical Computing, Vienna, Austria. URL http://www.R-project.org/).

39. Pritchard JK, Stephens M, Donnelly P (2000) Inference of population structure using multilocus genotype data. Genetics 155: 945–959.

40. Evanno, G., S. Regnaut, et al. (2005). “Detecting the number of clusters of individuals using the software STRUCTURE: a simulation study.” Mol Ecol 14(8): 2611–2620.

41. Earl, Dent A. and, vonHoldt, Bridgett M. (2012) STRUCTURE HARVESTER: a website and program for visualizing STRUCTURE output and implementing the Evanno method. Conservation Genetics Resources vol. 4 (2) pp. 359–361

42. Camacho C, Coulouris G, Avagyan V, Ma N, Papadopoulos J, et al. (2009) BLAST+: architecture and applications. BMC Bioinformatics 10: 421.

43. Huson DH, Bryant D (2006) Application of phylogenetic networks in evolutionary studies. Mol Biol Evol 23: 254–267.

44. Wagner CE, Keller I, Wittwer S, Selz OM, Mwaiko S, et al. (2013) Genome-wide RAD sequence data provide unprecedented resolution of species boundaries and relationships in the Lake Victoria cichlid adaptive radiation. Mol Ecol 22: 787–798.

45. Stamatakis A (2006) RAxML-VI-HPC: maximum likelihood-based phylogenetic analyses with thousands of taxa and mixed models. Bioinformatics 22: 2688–2690.

46. Heled J, Drummond AJ (2010) Bayesian inference of species trees from multilocus data. Mol Biol Evol 27: 570–580.

